# Inhibitory NPY neurons provide a large and heterotopic commissural projection in the inferior colliculus

**DOI:** 10.1101/2022.02.08.479582

**Authors:** Justin D. Anair, Marina A. Silveira, Pooyan Mirjalili, Nichole L. Beebe, Brett R. Schofield, Michael T. Roberts

## Abstract

Located in the midbrain, the inferior colliculus (IC) plays an essential role in many auditory computations, including speech processing and sound localization. The right and left side of the IC are interconnected by a dense fiber tract, the commissure of the IC (CoIC), that provides each IC with one its largest sources of input (i.e., the contralateral IC). Despite its prominence, the CoIC remains poorly understood. Previous studies using anterograde and retrograde tract-tracing showed that IC commissural projections are predominately homotopic and tonotopic, targeting mirror-image locations in the same frequency region in the contralateral IC. However, it is unknown whether specific classes of neurons, particularly inhibitory neurons which constitute ∼10-40% of the commissural projection, follow this pattern. We therefore examined the commissural projections of Neuropeptide Y (NPY) neurons, the first molecularly identifiable class of GABAergic neurons in the IC. Using retrograde tracing with Retrobeads in NPY-hrGFP mice of both sexes, we found that NPY neurons comprise ∼11% of the commissural projection. Moreover, focal injections of Retrobeads showed that NPY neurons in the central nucleus of the IC exhibit a more divergent and heterotopic commissural projection pattern than non-NPY neurons. Thus, commissural NPY neurons are positioned to provide cross-frequency, lateral inhibition to the contralateral IC. Through this circuit, sounds that drive activity in limited frequency bands on one side of the IC might suppress activity across a larger number of frequency bands in the contralateral IC.

## Introduction

Located in the midbrain, the inferior colliculus (IC) integrates inputs from the lower auditory brainstem and provides the major auditory projection to the thalamocortical system. Because of its strategic position, the IC is commonly described as the hub of the central auditory pathway (Adams, 1979; Cant and Benson, 2006). The IC is essential for most aspects of hearing, including speech computations, sound localization, and plasticity after hearing loss (Chase and Young, 2008; Carney et al., 2015; Chambers et al., 2016). Neurons in the IC are also important for binaural integration of sound (Litovsky et al., 2002; Sayegh et al., 2014), in which a key player is the commissure of the IC (CoIC), a large fiber tract that connects the two ICs and provides a substantial opportunity for integration of information across the midline (Aitkin and Phillips, 1984; González Hernández et al., 1986). The CoIC represents one of the largest pathways into or out of the IC (Moore, 1988), however the projection patterns and functional roles of individual classes of commissural neurons remain unknown.

Previous anatomical studies using anterograde and retrograde tracing found that commissural projections in the central nucleus of the IC (ICc) are largely homotopic and tonotopic, with commissural ICc neurons projecting to mirror-image frequency regions of the contralateral ICc (González-Hernández et al., 1986; Saldaña and Merchán, 1992; Malmierca et al., 1995, 2009). In contrast, neurons in the dorsal cortex of the IC (ICd) present two different projection patterns to the contralateral IC, with some neurons projecting to the ICd and others projecting to tonotopically matched regions of the ICc (Malmierca et al., 2009). Both GABAergic and glutamatergic neurons contribute to the CoIC, with inhibitory neurons comprising ∼10-40% of commissural projections (Malmierca et al., 1995; González-Hernández et al., 1996; Hernández et al., 2006; Nakamoto et al., 2013; Chen et al., 2018). Although inhibitory projections make up a minority of the commissural pathway, in vitro physiological studies using electrical stimulation (Smith, 1992; Moore et al., 1998; Li et al., 1999; Reetz and Ehret, 1999) and optogenetics (Goyer et al., 2019) report that, at the functional level, GABAergic synaptic inputs are more common than expected based on their anatomical percentage.

Functional studies indicate that commissural projections generally enhance the ability of neurons in the contralateral IC to detect and discriminate tones and sound localization cues by regulating the gain of auditory input-output functions (Malmierca et al., 2003, 2005; Orton and Rees, 2014; Orton et al., 2016). In individual neurons, commissural inputs could increase or decrease firing rates elicited by monaural or binaural sounds and could broaden or narrow frequency tuning curves. These results suggest that commissural influence can have net excitatory or inhibitory effects. While the commissural pathway itself contains excitatory and inhibitory projections, stimulation of the commissure activates feedforward circuits in the target IC that can drive secondary excitation and inhibition (Smith, 1992). It is not clear how inhibitory and excitatory commissural projections separately contribute to commissural computations.

The presence of a significant inhibitory component in the IC commissure along with the critical role of neural inhibition in shaping auditory cues in the IC (Pollak et al., 2011) raises the question of how GABAergic neurons contribute to commissural function. However, the diversity of GABAergic neurons in the IC (Ono et al., 2005a; Beebe et al., 2016) raises the possibility that different classes of GABAergic neurons play distinct roles in commissural modulation of sound processing. As a step towards understanding the organization of inhibitory neural circuits in the IC, we recently identified Neuropeptide Y (NPY) neurons as the first molecularly identifiable class of GABAergic neurons in the IC (Silveira et al., 2020). NPY neurons are labeled in NPY-hrGFP mice (van den Pol et al., 2009) and are principal neurons that send long-range inhibitory projections to the auditory thalamus. NPY neurons have a stellate morphology and represent approximately one-third of IC GABAergic neurons (Silveira et al., 2020).

Here, using retrograde tracing with red Retrobeads (RB) in NPY-hrGFP mice, we found that NPY neurons project to the contralateral IC. Although only a small proportion of NPY neurons contributed to the CoIC projection, NPY neurons comprised ∼11% of commissural neurons, representing a large portion of the inhibitory commissural projection. Focal injections of RB in the ICc showed that non-NPY commissural neurons tended to target homotopic frequency bands in the contralateral ICc while NPY neurons had a more divergent and heterotopic projection pattern, mostly targeting frequency regions bordering the main frequency band labeled by non-NPY neurons. NPY neurons also participated in the commissural projection between the left and right ICd, but in the ICd the NPY projection was organized similarly to the non-NPY projection. These results provide the first insights into the organization and possible functional roles of the inhibitory commissural projection. Based on our results, we propose that commissural NPY neurons provide cross-frequency lateral inhibition to the contralateral ICc and therefore may constitute an important contralateral source for sideband inhibition.

## Materials and Methods

### Animals

All experiments were approved by the University of Michigan Institutional Animal Care and Use Committee and followed NIH guidelines for the use and care of laboratory animals. All animals were kept on a 12-hour day/night cycle with unrestricted access to food and water. To visualize NPY neurons, NPY-hrGFP mice were obtained from Jackson Laboratory (stock #006417) and were kept hemizygous for the hrGFP transgene by crossing with C57BL/6J mice (Jackson Laboratory; stock #000664)(van den Pol et al., 2009). Mice used for experiments were aged P22-P69 to avoid possible changes to auditory structures as a result from age-related hearing loss due to the *Cdh23*^*ahl*^ mutation present in C57BL/6J mice (Noben-Trauth et al., 2003). For all experiments, mice of both sexes were used.

### Intracranial Retrobead Injections

To quantify and determine the distribution of CoIC projections of NPY neurons we performed retrograde tracing using red fluorescent Retrobeads (RB, “red beads,” Luma-Fluor, Inc., Naples, Fl, USA; 1:2 - 1:4 dilution). Injections were made into the right IC of 10 NPY-hrGFP mice aged P22-P62 of both sexes. Throughout the procedure, mice were anesthetized with isoflurane (1.5-3%) and their body temperature maintained with a homeothermic heating pad. Mice were injected subcutaneously with the analgesic carprofen (5mg/kg, CarproJect, Henry Schein Animal Health). Following this, the scalp was shaved, and a rostro-caudal incision was made to expose the skull. The injection sites were mapped using stereotaxic coordinates relative to the lambda suture, and injection depths were relative to the surface of the skull. A single craniotomy was made above the injection site using a micromotor drill (K 1050, Foredom Electric Co.) with a 0.5 mm burr (Fine Science Tools). For the first set of experiments, we made multi-subdivision RB injections into the right IC through two penetrations: penetration 1 – 900 µm caudal, 1000 µm lateral, and 2000 µm deep; penetration 2 – 900 µm caudal, 1250 µm lateral, and 2000 µm deep. One injection of 40 - 60 nl of RB was performed in each penetration, resulting in two RB deposits with a total volume of 80 - 120 nl. For the second set of experiments, we made focal injections in one penetration: -900 μm caudal, 1000-1150 μm lateral and 1600-1750 μm deep for the ICc injections and - 900 μm caudal, 875 μm lateral, and 1775 μm deep for the ICd injection.

RB were injected with a Nanoject III nanoliter injector (Drummond Scientific Company) connected to an MP-285 micromanipulator (Sutter Instruments). Glass injection pipettes were pulled from 1.14 mm outer diameter, 0.53 mm inner diameter capillary glass (cat# 3-000-203-G/X, Drummond Scientific Company) with a P-1000 microelectrode puller (Sutter Instrument). The injector tip was cut, and front filled with RB. After the injection was complete, the scalp was sutured using Ethicon 6-0 (0.7 metric) nylon sutures (Ethicon USA LLC) or glued using 3M Vetbond tissue adhesive (3M), and the incision was treated with 0.5 mL 2% lidocaine hydrochloride jelly (Akorn Inc.). After recovery from anesthesia, mice were returned to the vivarium and were monitored daily.

### Immunofluorescence

Five to ten days after RB injection, mice were anesthetized with isoflurane and perfused transcardially with 0.1 M phosphate-buffered saline (PBS), pH = 7.4, for 1 min and then with a 10% buffered formalin solution (Millipore Sigma, cat# HT501128) for 10 min. Brains were collected and stored in the same formalin solution for 2 hours then cryoprotected overnight at 4°C in 0.1 M PBS containing 20% sucrose. Brains were cut into 40 - 50 μm coronal sections on a vibratome or freezing microtome. Sections were washed in PBS (3 washes of 10 minutes each), and then treated with 10% normal donkey serum (Jackson ImmunoResearch Laboratories, catalog #017-000-121) and 0.3% Triton X-100 in PBS for 2 h. Sections were then incubated for 24-40h at 4°C in mouse anti-GAD67 (1:1000; Sigma Millipore, catalog #MAB5406), rabbit anti-NeuN (1:500; Sigma Millipore, catalog #ABN78), and/or guinea pig anti-GlyT2 (1:2000; Synaptic Systems, cata-log #272-004). After the incubation period, sections were rinsed in PBS and incubated in AlexaFluor-647-tagged goat anti-mouse IgG (1:100; Thermo Fisher Scientific, catalog #A21039), AlexaFluor-647-tagged donkey anti-mouse IgG, AlexaFluor-647-tagged donkey anti-rabbit IgG, and/or AlexaFluor-594-tagged goat anti-guinea pig IgG (1:500; Thermo Fisher Scientific, catalog #A-21202, #A-21206, and #A-11076) for 1.5-2h at room temperature. Sections were then mounted on Superfrost Plus microscope slides (Thermo Fisher Scientific, catalog #12-550-15) and coverslipped using Fluoromount-G (SouthernBiotech, catalog #0100–01) or DPX (Sigma Millipore, catalog #06522). For the multi-subdivision RB injections, images were collected using a Zeiss AxioImager.Z2 microscope. Low magnification images were collected using a 5X objective. High-magnification images are maximum intensity projections of image stacks obtained with a 63X oil-immersion objective (NA = 1.4) and structured illumination (Apotome 2, Zeiss) to provide optical sectioning at 0.5 µm z-steps. For the focal RB injections, images of the IC were collected using a 10x objective, 20x objective, 1.30 NA 40x oil-immersion objective (1024 × 1024 resolution, 1 μm z-step, 0.75 zoom), or a 1.40 NA 63x oil-immersion objective (1024 × 1024 resolution, 1 μm z-step, 0.75 zoom) on a Leica TCS SP8 laser scanning confocal microscope.

### Antibody Characterization

GABAergic neurons were identified using the mouse monoclonal anti-GAD67 antibody (Sigma Millipore, catalog #MAB5406). The 67 kDA isoform of glutamic acid-decarboxylase (GAD) is required for the synthesis of the neurotransmitter GABA. Anti-GAD67 antibodies were raised against this isoform. The vendor reported no cross-reactivity for GAD65 – the 65 kDA isoform of GAD – using a Western blot analysis. The same anti-GAD67 antibody has been used in several IC studies to identify GABAergic neurons (Ito et al., 2009; Beebe et al., 2016; Goyer et al., 2019; Silveira et al., 2020). To label glycinergic terminals, we used a guinea pig polyclonal anti-GlyT2 antibody (Synaptic Systems, catalog #272004). Glycine transporter 2 (Glyt2) is a transmembrane protein involved in the removal of extracellular glycine and primarily labels axons and terminals. The manufacturer reported complete specificity for the 100 kDA isoform of GlyT2 using Western blot analysis in mouse brain tissue. To visualize neuronal cell bodies, we performed anti-NeuN staining with a rabbit polyclonal antibody (Sigma Millipore, catalog #ABN78). Using a Western blot analysis on mouse brain tissue, this antibody showed to selectively bind a set of closely associated protein isoforms of NeuN, a neuron specific protein which binds to DNA in neurons. Several studies have used this antibody as a general neuronal marker in the IC (Mellott et al., 2014; Beebe et al., 2016; Goyer et al., 2019; Silveira et al., 2020; Beebe and Schofield, 2021).

### IC Subdivision Delineation

Due to fluorescence overlap, immunofluorescence to delineate IC subdivisions was not performed in sections from mice injected with RB. Instead, we prepared a reference series of sections from a control C57BL/6J mouse, aged P49. These sections were immunolabeled for GAD67 and GlyT2 and imaged using the 20x objective of a Leica TCS SP8 confocal microscope. The pattern of GAD67 and GlyT2 immunolabeling has previously been used to identify IC subdivisions (Buentello et al., 2015; Silveira et al., 2020; Beebe and Schofield, 2021). For each RB injection case, IC subdivision boundaries were drawn based on the distribution of GABA and glycinergic terminals in the most similar reference sections. The outer edges of the IC were identified based on fluorescence from the NeuN immunolabeling. The outline and subdivision borders of the IC were drawn using Neurolucida 11 software (MBF Bioscience, Williston, VT, USA).

### Retrobead Analysis

Following multi-subdivision RB injections, a series of every third section through the rostro-caudal extent of the contralateral IC in each case was examined for hrGFP^+^ (NPY) cells, and for RB-labeled cells (which were presumed to project across the CoIC). Cells of either type, or those where both markers were present, were plotted using a Zeiss AxioImager.Z2 microscope and Neurolucida 11 software (MBF Bioscience). Marker counts and plots were exported from Neurolucida Explorer and prepared using Microsoft Excel and Adobe Illustrator, respectively.

Following focal RB injections, confocal images were taken from a series of 5-8 coronal sections evenly spaced along the rostral-caudal extent of the IC and imported into Neurolucida 11 software (MBF Bioscience) for analysis. NPY neurons were identified based on hrGFP fluorescence and marked when RB clearly filled and co-labeled an hrGFP^+^ cell body. Similarly, non-NPY neurons were identified and marked when RB clearly filled and co-labeled a NeuN+/hrGFP-negative cell body. Contours and plots of the left IC (contralateral to the RB injection site) were exported from Neurolucida 11 and assembled into figures in Adobe Illustrator 2022.

### Cumulative Distribution Analysis

For cases where the RB injection was centered in the ICc, the analyzed sections were imported into the serial section manager of Neurolucida 11. The sections were oriented and stacked in 3D from caudal (bottom) to rostral (top). To assess if RB-labeled NPY neurons had a different distribution than the RB-labeled non-NPY neurons, we compared the distances of each RB-labeled neuron to a point, which we refer to as the centroid, located approximately homotopic to the center of mass of the RB injection site in the contralateral IC. To define the centroid point, we first collected a 10x image of the IC slice containing the strongest RB labeling on the injected side (the right side) of the IC. We then imported this image to Neurolucida and used contours to outline the perimeter of the IC and the main body of the RB labeling corresponding to the injection site. These contours were then reflected across the midline of the IC to yield a “mirror-image” projection of the injection site, which was aligned to the previously prepared reconstruction of the left IC. To prevent bias in this alignment procedure, cell markers were hidden so that only contours were visible. The coordinates of the centroid of the mirror-image injection site were calculated using the Contour Analysis function in Neurolucida Explorer (MBF Bioscience). A locus marker was placed at the centroid coordinates, and the distances, in µm, between each neuron marker and the centroid were calculated using the Marker Analysis functions in Neurolucida Explorer. These distances were imported to Igor Pro 9 (WaveMetrics), and for each case, normalized cumulative distribution plots with the same number of bins, bin width = 20 µm, were constructed for both NPY and non-NPY neuron populations.

### Heatmap Analysis

Serial sections were aligned and stacked in Neurolucida as described above for the Cumulative Distribution Analysis. *X* and *Y* coordinates from each neuron were then extracted in Neurolucida Explorer and imported into MATLAB (MathWorks). In MATLAB, the numbers of neurons in 100 µm x 100 µm bins covering the entire extent of the left IC were counted using the neuron coordinates and the “hist3” function. The resulting bivariate histograms were plotted as heatmaps where the intensity of the color corresponds to the densities of NPY and non-NPY neurons. Overlays of the NPY and non-NPY neuron heatmaps were prepared using the “imfuse” function in MATLAB.

## Results

To compare the commissural projection patterns of NPY neurons and non-NPY neurons, injections of RB were made in the right IC of NPY-hrGFP mice of both sexes. Multi-subdivision RB injections were made in 4 mice to assess the overall distribution of NPY neurons contributing to the commissural pathway. Focal RB injections were made in 6 mice to assess and compare the topography of NPY and non-NPY commissural projections.

### NPY neurons provide a large portion of the inhibitory commissural projection

Multi-subdivision injections included two or more IC subdivisions and extended across more of the IC rostro-caudal axis as compared to focal injections. **Figure 1A** shows an injection that included lateral parts of the ICd and dorsal parts of the ICc and IClc. Other cases included more ventral parts of the ICc and IClc. These multi-subdivision injections typically resulted in many RB-labeled neurons in auditory centers, including the contralateral IC. RB-labeled cells were present widely across the subdivisions of the contralateral IC and could be NPY+ or NPY-. **Figure 1B** shows examples of RB-labeled cells in each subdivision of the contralateral IC, some of which are NPY+ (magenta arrows), and some of which are NPY-negative (“non-NPY”; cyan arrowheads). NPY+ cells that lacked retrograde tracer were also observed in the contralateral IC (green arrowheads). **Figure 1C** shows the distribution of RB-labeled commissural cells, including non-NPY cells (cyan squares) and NPY+ cells (magenta triangles) in the IC contralateral to an RB injection in one case. Across four cases with multi-subdivision injections, an average of 11.1% ± 2.1% of CoIC-projecting cells were NPY+, with similar proportions (10-12%) in individual IC subdivisions (**Figure 1D**). This suggests that NPY cells make up a substantial portion, potentially a majority, of inhibitory commissural cells, which have been estimated to be 10-40% of the overall CoIC pathway in other studies (Malmierca et al., 1995; González-Hernández et al., 1996; Nakamoto et al., 2013; Chen et al., 2018).

**Figure 1.**
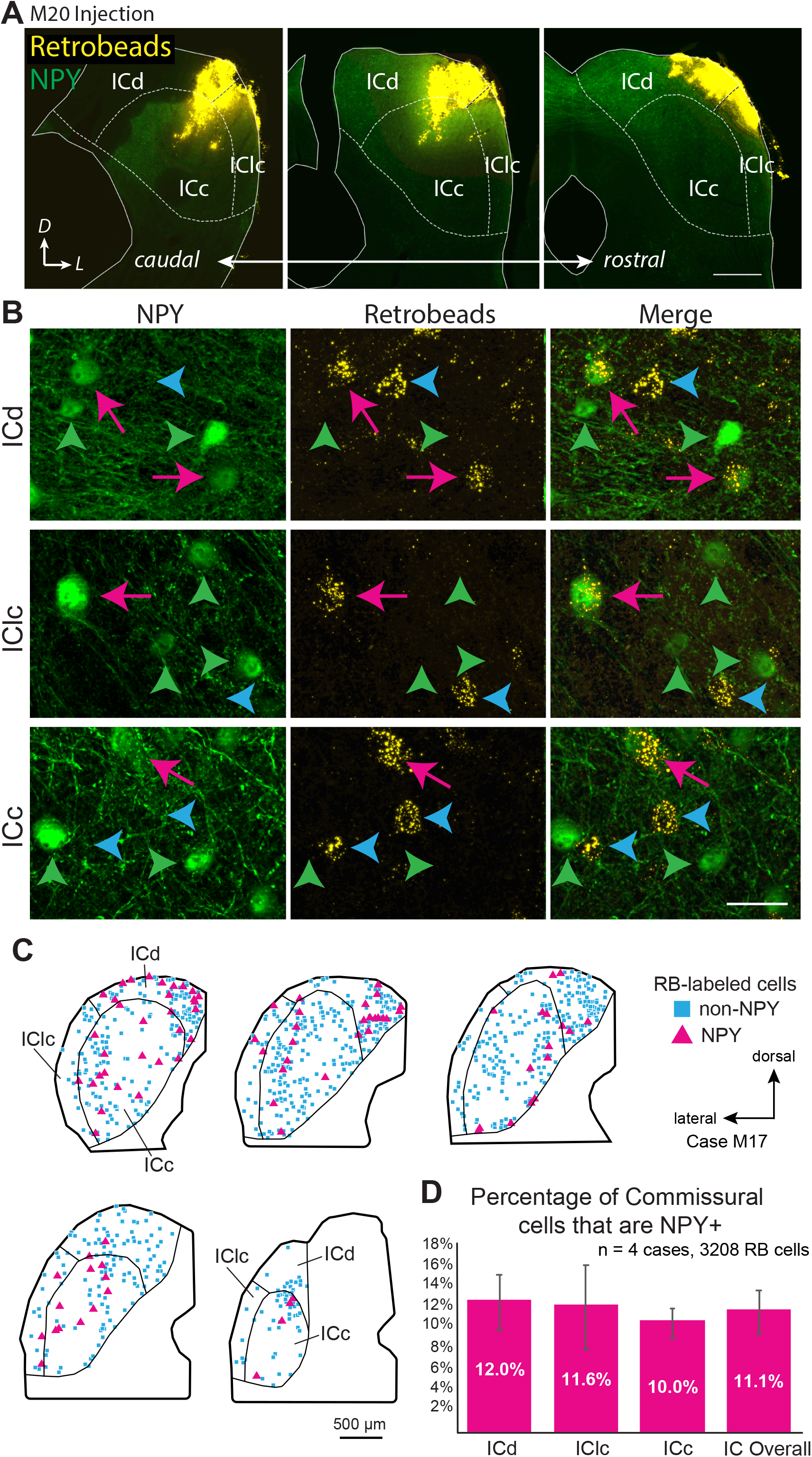
Deposits of retrograde tracer across multiple IC subdivisions labels NPY+ and NPY-negative (“non-NPY”) commissural cells in all subdivisions of the contralateral IC. **A)** Three images in a caudal to rostral sequence showing a deposit of Retrobeads (RB, yellow) in the right IC that included parts of the central IC (ICc), dorsal cortex (ICd) and lateral cortex (IClc). The injection was made in an NPY-hrGFP mouse so NPY is green. Scale bar = 0.5 mm. **B)** High magnification images show cells in the left IC that were labeled by an injection of RB in the right IC. Examples are shown from the dorsal cortex (ICd, top row), lateral cortex (IClc, middle row) and central nucleus (ICc, bottom row). In each row, the first column shows NPY-hrGFP label and the second column shows RB labeling in the same field of view (merged view in third column). RB-labeled cells could be double labeled with NPY (magenta arrows) or could be NPY-negative (“non-NPY”, cyan arrowheads). Also present are NPY+ cells that did not contain RB (green arrowheads). Scale bar = 25 µm. **C)** Plots showing the distribution of RB-labeled commissural cells in a series of transverse sections. A majority of RB-labeled neurons were non-NPY (cyan); NPY+ commissural cells (magenta) were interspersed among the non-NPY cells throughout the IC. **D)** Graph showing the average percentage of RB-labeled (i.e., commissural) cells that were NPY+ in the IC overall and in each of the major subdivisions across four cases. NPY cells contribute roughly 10-12% of the commissural projection. Error bars = SD.

### Commissural projections from ICc NPY neurons are more divergent, less homotopic, than those of non-NPY projections

Previous studies showed that the commissural projections of ICc neurons are predominately homotopic and tonotopic, connecting similarly tuned isofrequency lamina between the two sides of the IC (Saldaña and Merchán, 1992; Malmierca et al., 1995, 2009). However, these studies did not distinguish among neuron types, and since GABAergic neurons comprise only ∼25% of IC neurons and ∼10-40% of commissural neurons, it is likely that past results mainly reflect the projection patterns of excitatory IC neurons. To test whether NPY neurons follow the projection pattern previously described, we made small, focal injections of RB in the right ICc of 5 NPY-hrGFP mice. Following each injection, the left IC contained NPY neurons and non-NPY neurons labeled with RB, indicating that they formed commissural projections to the injection site in the right IC (**Figure 2**).

**Figure 2.**
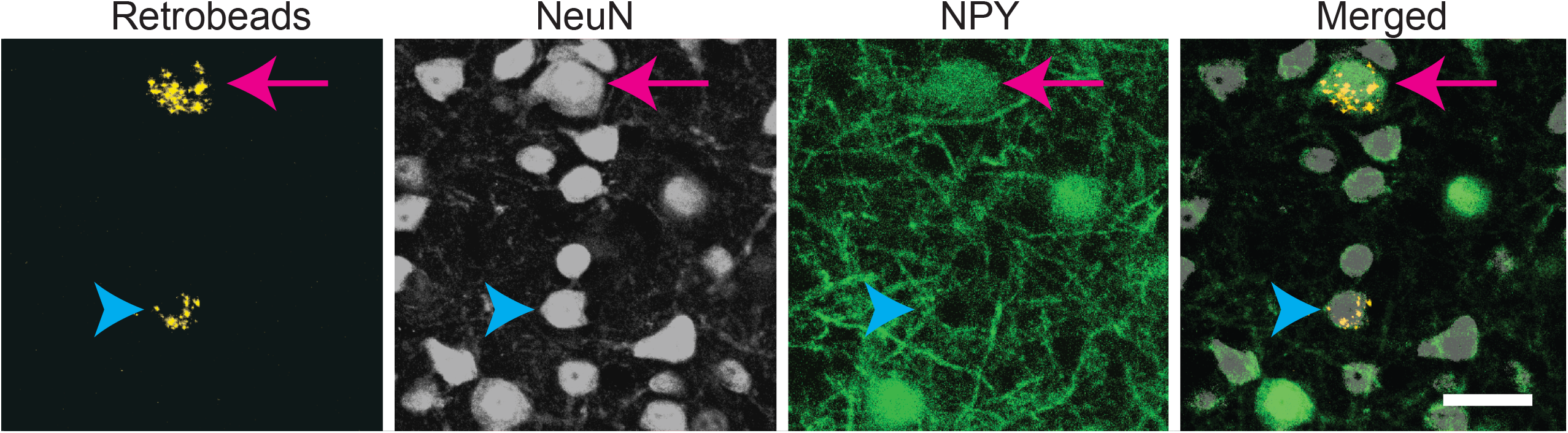
Both NPY+ and non-NPY neurons were labeled by Retrobeads after tracer deposits restricted to a single IC subdivision. High magnification images from the left ICc following a 10 nl injection of RB into the right ICc of an NPY-hrGFP mouse. An NPY+ neuron (arrow) and a non-NPY neuron (arrowhead) co-labeled with RB, indicating that they projected to the contralateral ICc. Scale bar = 25 µm.

For each ICc injection case, we analyzed the location of the RB injection site in the right IC and the distribution of RB-labeled neurons in the left IC. In Case 25, nearly all the RB deposit was in a narrow tract in the dorsal half of the right ICc (**Figure 3A**). In the left IC, non-NPY neurons labeled with RB formed a densely packed cluster that extended throughout the ICc along a ventrolateral to dorsomedial axis, with relatively narrow spread perpendicular to this main axis. Fewer RB-labeled non-NPY neurons were present in the ICd and very few were observed in the IClc (**Figure 3B**, cyan squares). This pattern remained similar across much of the rostro-caudal extent of the ICc but tapered off in more rostral sections. RB-labeling was particularly dense at the location mirroring the injection site in the right IC (**Figure 3B**, gold outline in section 3). These results are consistent with the commissural projection patterns detailed in previous studies, suggesting that the commissural projections of non-NPY neurons mainly connect corresponding isofrequency lamina on the two sides of the ICc. In contrast, RB-labeled NPY neurons within the ICc were more common on the outer edges of the region defined by the RB-labeled, non-NPY neurons.

**Figure 3.**
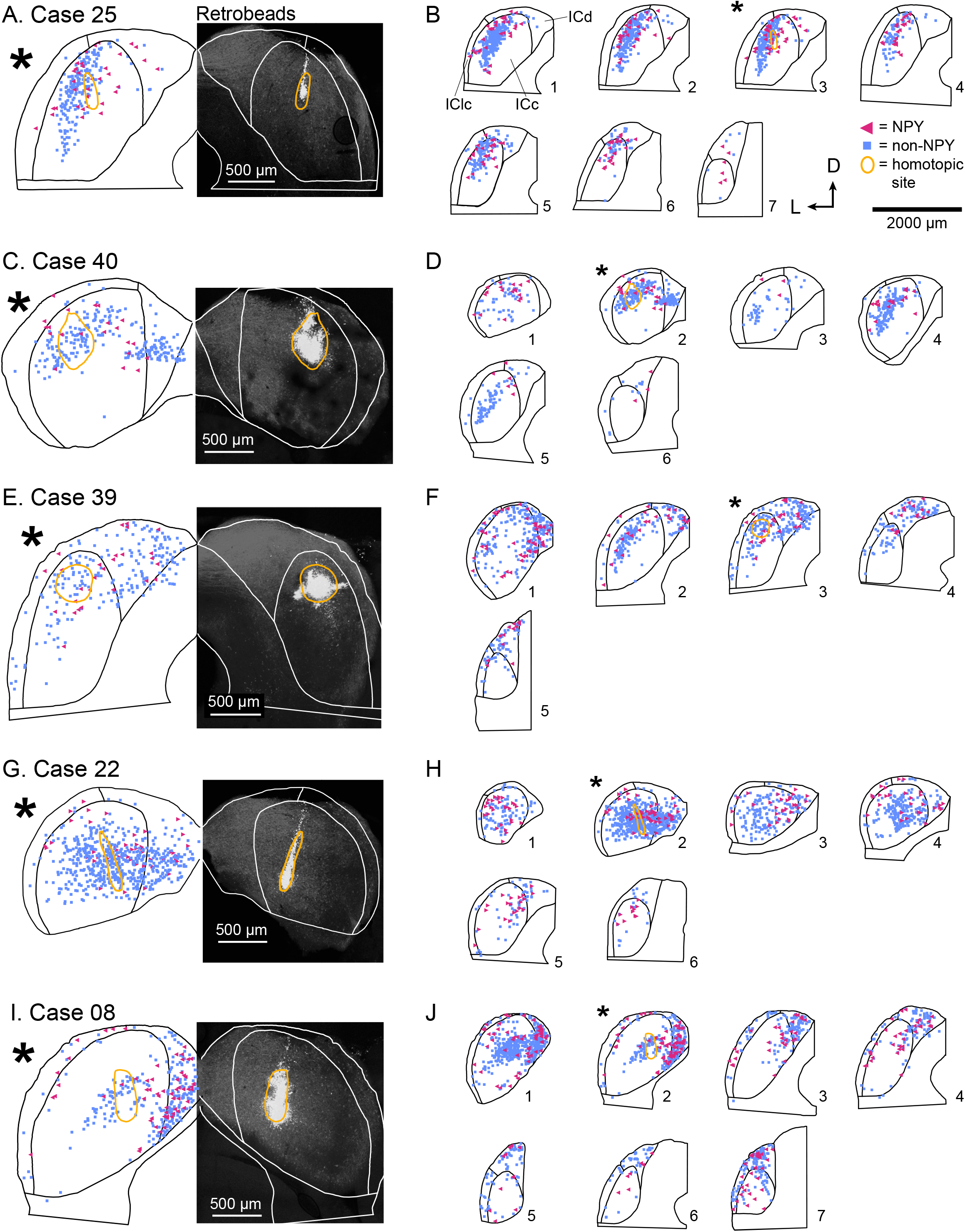
In the ICc, commissural projections of NPY neurons are more heterotopic than those of non-NPY neurons. The panel shows results from five cases where injections were predominately limited to the ICc. **A)** RB deposit site in the right IC and homotopic site in the left IC in experimental Case 25. The right half of the panel includes a gray-scale image showing the center of the RB deposit (circled in gold). The left half of the panel shows an outline of the left IC with the homotopic location of the injection site indicated by a reflected image of the gold outline. The illustrated section (*) is from a series of sections shown in **B. B)** The full series of plots for Case 25 showing the distributions of RB-labeled (commissural) neurons that were NPY+ (magenta triangles) or non-NPY (cyan squares). Sections are numbered from caudal to rostral in each series. The asterisk indicates the section shown in **A** with the area homotopic to the injection outlined in gold. D – dorsal; L – lateral. **C-J)** Results from four additional experimental cases, displayed as in **A** and **B**. The locations of the RB deposits in the ICc varied across cases, but the differences in distributions of NPY vs. non-NPY commissural cells was consistent: the NPY cells were less focused in the homotopic site. The main pattern of non-NPY neuron labeling in the ICc was often consistent with the shape of a frequency band, while NPY+ neurons were more common in regions bordering the frequency band. **B**, *n* = 898 non-NPY neurons, 168 NPY neurons. **D**, *n* = 573 non-NPY neurons, 86 NPY neurons. **F**, *n* = 1095 non-NPY neurons, 161 NPY neurons. **H**, *n* = 782 non-NPY neurons, 135 NPY neurons. **J**, *n* = 1173 non-NPY neurons, 226 NPY neurons.

In Case 40, the RB injection site was in the dorsal half of the right ICc and was more caudal than the injection site for Case 25 (**Figure 3C**). RB-labeled NPY and non-NPY neurons were present in the left ICc and ICd, with little labeling in the IClc (**Figure 3D**). As with Case 25, RB-labeling of non-NPY neurons in the left ICc was dense at the site homotopic to the injection site (**Figure 3D**, gold outline in section 2) and mainly extended in a band matching the presumed orientation of the isofrequency laminae (**Figure 3D**, cyan squares). The distribution of RB-labeled NPY neurons partly overlapped the distribution of non-NPY neurons, but also appeared more dispersed (**Figure 3D**, magenta triangles).

In Case 39, the RB deposit was similar in location but larger than that in Case 25, located in the dorsal half of the right ICc, slightly rostral to the midpoint of the rostral-caudal axis of the IC (**Figure 3E**). RB-labeling of non-NPY neurons in the left ICc was strongest around the site homotopic to the injection site and extended through the ICc in a pattern consistent with labeling of a small number of isofrequency laminae (**Figure 3F**). In contrast, RB-labeled NPY neurons again appeared more common in ICc regions surrounding the region where RB-labeled non-NPY neurons were densest (**Figure 3F**). RB-labeled NPY and non-NPY neurons were also common in the ICd and rarer in the IClc, although there were more RB-labeled neurons in the IClc in this case than the previous cases. Despite the variation in the size of tracer deposit and its rostro-caudal location, the results from the three cases with injections in the dorsal ICc showed similar results.

In Case 22, the RB injection site included some dorsal ICc but extended will into the ventral ICc, producing a deposit much more spread out than in the previous cases (**Figure 3G**). In the left IC, RB-labeled NPY and non-NPY neurons were again common in the ICc and ICd, but rare in the IClc. Within the ICc, the labeling of non-NPY neurons did not produce a distribution as narrow as the previous cases. Nonetheless, the distribution of RB-labeled NPY neurons in the left IC appeared more dispersed and less prevalent at the site homotopic to the RB injection site compared to non-NPY neurons (**Figure 3H**).

In Case 08, the final ICc injection case, the RB injection site was in a caudal and ventral portion of the ICc, closer to the midline than in the previous cases (**Figure 3I**). RB-labeled non-NPY neurons were dense at the site homotopic to the injection site and again extended out from the homotopic site following a laminar pattern that extended through the ICc (**Figure 3J**, sections 1 and 2). Once again, RB-labeled NPY neurons were more common in the regions surrounding the homotopic site (**Figure 3J**).

#### Heatmap analysis

To summarize the distribution of RB-labeled neurons in the ICc across coronal sections, we aligned IC sections according to the three-dimensional shape of the IC, compressed neuron distributions into two dimensions, removed markers for neurons not located in the ICc, and generated heatmaps to separately show the density distributions of NPY and non-NPY neurons in the ICc (**Figure 4**). For Case 25, the heatmaps show that the densest commissural projection of non-NPY neurons was from a site homotopic to the RB injection site (**Figure 4A, left**). The remainder of the non-NPY neurons projection came from an angled tract consistent with the presumed orientation of the isofrequency laminae, with the density of labeled cells decreasing with increasing distance from the homotopic site. In contrast, the distribution of RB-labeled NPY neurons was densest at sites surrounding the homotopic site, with a comparatively low density at the homotopic site (**Figure 4A**, middle). An overlay of the non-NPY and NPY heatmaps shows that the NPY neuron distribution was broader than the non-NPY distribution along the dorsolateral-ventromedial axis (i.e., perpendicular to the presumed isofrequency axis) (**Figure 4A, right**). The two cell populations spanned a similar distance along the isofrequency axis. A similar pattern was apparent in the heatmaps for Case 40, which again showed that RB-labeled non-NPY neurons tended to follow a laminar distribution (**Figure 4B**), while RB-labeled NPY neurons were more dispersed, with highest densities in regions away from the peak densities of non-NPY neurons.

**Figure 4.**
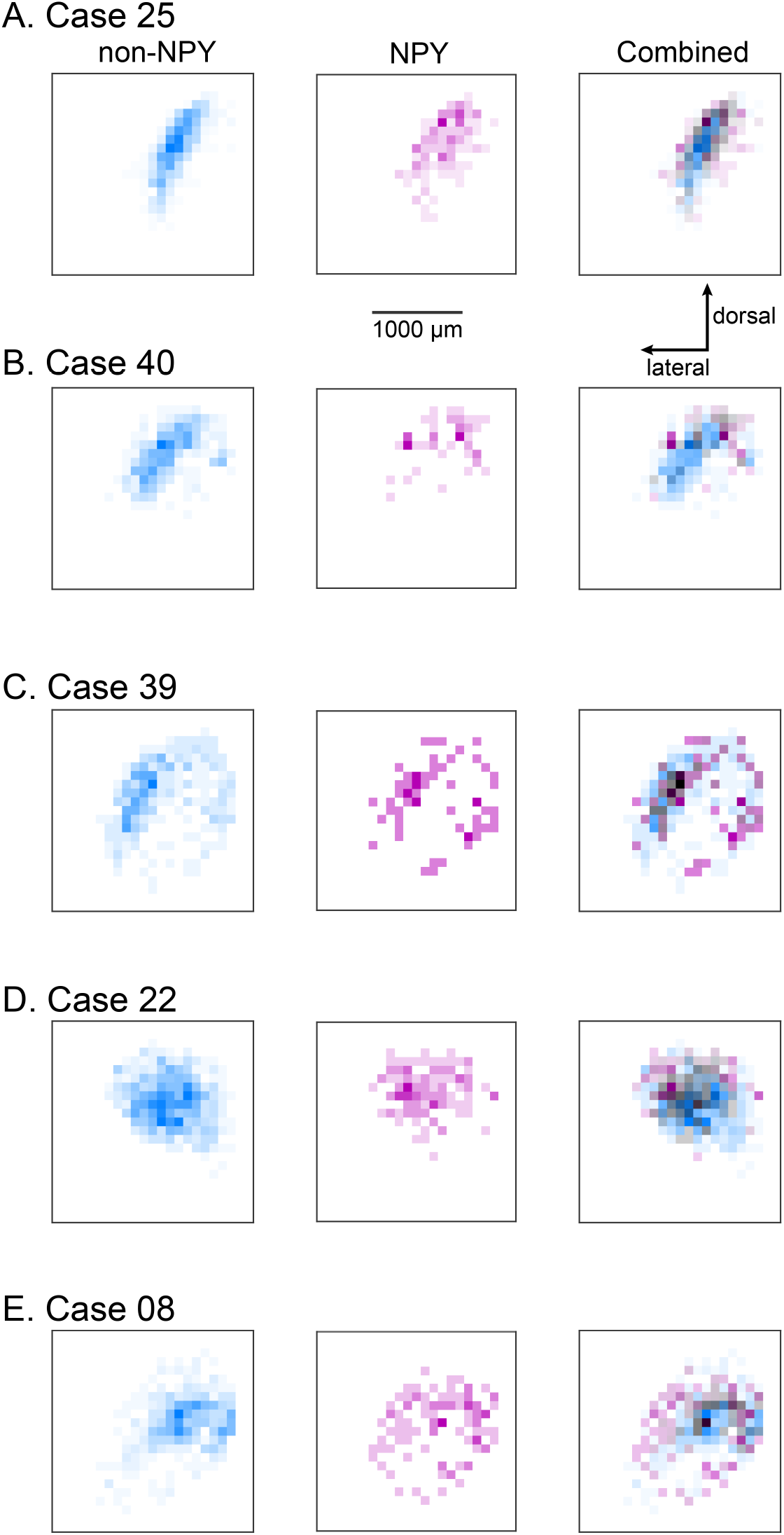
Heatmaps highlight the more heterotopic distribution of RB-labeled NPY neurons in the ICc compared to RB-labeled non-NPY neurons. For each of the ICc injection cases, reconstructions of coronal sections of the left IC were aligned and flattened in the rostral-caudal dimension (Z-axis), and the density of RB-labeled neurons in the ICc was calculated in the coronal plane using 100 µm^2^ bins. RB-labeled neurons in the ICd and IClc were excluded from this analysis. **A-E)** The density of RB-labeled non-NPY neurons in the left ICc is shown in cyan (left), while the density of RB-labeled NPY neurons in the left ICc is shown in magenta (middle). More intense colors indicate higher neuron densities normalized separately to the maximum density of non-NPY neurons (left) and NPY neurons (middle). Merged overlays of the non-NPY and NPY densities (right) reveal that NPY neurons were more common at sites surrounding the main mass of the non-NPY density. Bins where non-NPY and NPY neurons strongly overlapped appear in gray, with darker grays indicating more overlap. Case numbers are indicated in the subpanel labels. The scale bar and dorsal and lateral direction arrows in **A** also apply to **B-E**. In **A**,**B**,**C**, the main pattern of non-NPY neuron labeling follows a ventrolateral to dorsomedial orientation, consistent with labeling of one or more isofrequency laminae, and the distribution of NPY neurons in these cases generally shows higher densities at regions bordering the main band of non-NPY labeling. In **D**,**E**, the distribution of RB-labeled non-NPY neurons is broader, with a less laminar shape than observed in **A**,**B**,**C**, but RB-labeled NPY neurons remain more common at sites bordering the main pattern of non-NPY labeling.

In Cases 39, 22, and 08, heatmaps again showed that non-NPY neurons were densest in regions that were mainly homotopic to the RB injection site, while NPY neurons were denser in regions surrounding the area where non-NPY neurons were densest (**Figure 4C-E**). This trend was apparent even though the distribution of non-NPY neurons was less obviously following the arrangement of the isofrequency laminae in these cases. Together, heatmap analysis of the ICc injection cases suggests that the commissural projections of NPY neurons are more divergent than those of non-NPY neurons. Furthermore, given the tonotopic organization of the ICc, the larger spread of NPY neurons in the dorsolateral-ventromedial axis, corresponding to spread across more isofrequency laminae, suggests that commissural NPY neurons provide cross-frequency inhibition to neurons in the contralateral IC.

#### Quantitative analysis

To quantitatively assess the tendencies of NPY and non-NPY neurons in the ICc to make homotopic commissural projections, we mapped for each ICc case a point in the left IC homotopic to the centroid (center of mass) of the RB injection site in the right IC and measured the three-dimensional distance of each RB-labeled ICc neuron to this centroid. Cumulative probability plots show that NPY neurons in the ICc tended to be located farther from the homotopic centroid than non-NPY neurons in the ICc in all five ICc injection cases (**Figure 5A-E**). Two-sample Kolmogorov-Smirnov tests confirmed that the distributions of the distances of NPY and non-NPY neurons from the homotopic centroid were significantly different (p < 0.05) in each ICc case (Case 25: *n* = 832 non-NPY, 149 NPY, *D*_(832,149)_ = 0.15, *p* = 0.006; Case 40: *n* = 469 non-NPY, 61 NPY, *D*_(469,61)_ = 0.24, *p* = 0.003; Case 22: *n* = 872 non-NPY, 125 NPY, *D*_(872,125)_ = 0.20, *p* = 2e-4; Case 39: *n* = 387 non-NPY, 60 NPY, *D*_(387,60)_ = 0.21, *p* = 0.017; Case 08: *n* = 586 non-NPY, 104 NPY, *D*_(586,104)_ = 0.27, *p* = 3e-6). In addition, comparison of the medians of the distances from the homotopic centroid showed that NPY neurons were located significantly farther from the centroid than non-NPY neurons (**Figure 5F**; mean ± SD, non-NPY = 370 ± 61 µm, NPY = 458 ± 75 µm; paired t-test, *n* = 5, *t*_(4)_ = -8.23, *p* = 0.001).

**Figure 5.**
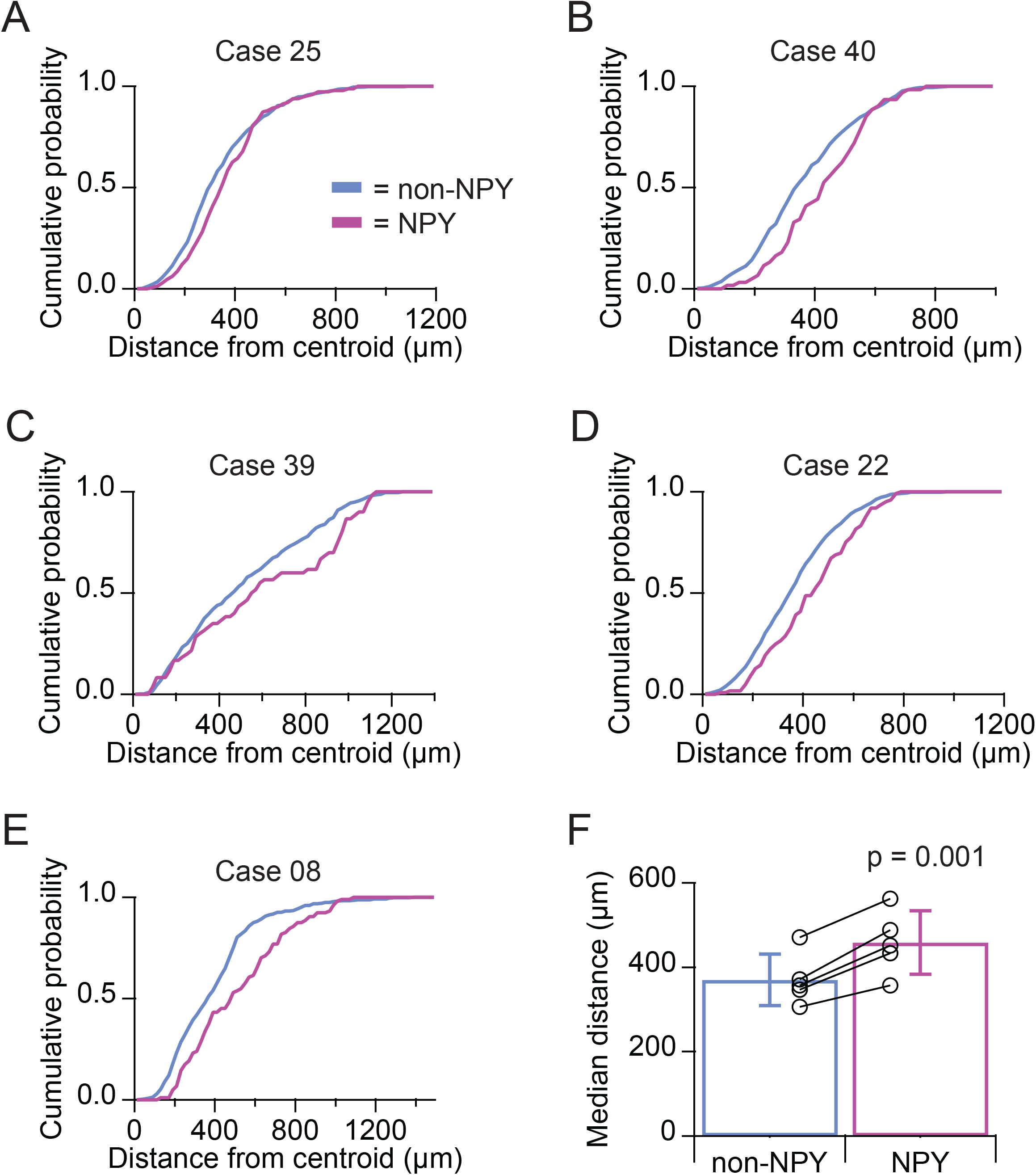
Quantitative analysis confirms that the commissural projections of NPY neurons in the ICc are more heterotopic than those of non-NPY neurons. For each of the ICc injection cases, reconstructions of coronal sections of the left IC were aligned, the site in the left ICc homotopic to the centroid of the injection site in the right IC was mapped, and the three-dimensional distances of RB-labeled non-NPY and NPY neurons to the homotopic centroid were measured. RB-labeled neurons in the ICd and IClc were excluded from this analysis. **A-E)** Cumulative distribution plots show that RB-labeled NPY neurons (magenta) tended to be located farther from the homotopic centroid than RB-labeled non-NPY neurons (cyan). In each case, a two-sample Kolmogorov-Smirnov test revealed that the distributions of distances to the centroid were significantly different for NPY and non-NPY neurons (see Results for details). **F)** Comparison of the median distances to the homotopic centroid from each ICc injection case also showed that NPY neurons were located significantly farther from the homotopic centroid than non-NPY neurons (paired t-test, see Results for details). Bars indicate mean ± SD of the median distances.

Since the preparation of serial brain sections can cause the measurement of distances across subsequent sections to be less accurate than the measurement of distances within sections, we repeated the above quantitative analyses on a flattened data set where the *Z*-dimension (rostral-caudal axis) was removed. Distributions of the two-dimensional distances of NPY and non-NPY ICc neurons from the homotopic centroid remained significantly different (p < 0.05) in each ICc case, with the distribution of NPY neurons shifted to the right indicating greater distances from the centroid (data not shown; Case 25: *D*_(832,149)_ = 0.16, *p* = 0.004; Case 40: *D*_(469,61)_ = 0.26, *p* = 0.001; Case 22: *D*_(872,125)_ = 0.14, *p* = 0.023; Case 39: *D*_(387,60)_ = 0.21, *p* = 0.015; Case 08: *D*_(586,104)_ = 0.26, *p* = 1.4e-5). Similarly, comparisons of the median two-dimensional distances of NPY and non-NPY neurons to the homotopic centroid again showed that NPY neurons were located significantly farther from the centroid (data not shown; mean ± SD, non-NPY = 334 ± 64 µm, NPY = 421 ± 81 µm; paired t-test, *n* = 5, *t*_(4)_ = -5.21, *p* = 0.006).

Combined, results from the five ICc injection cases indicate that the commissural projections of non-NPY neurons in the ICc are predominately homotopic and tonotopic, often following the laminar topology of the ICc. In contrast, the commissural projections of NPY neurons in the ICc were more heterotopic, commonly targeting contralateral sites that likely had somewhat higher or lower frequency tuning than the site where the projection originated. Thus, non-NPY commissural neurons in the ICc may predominately provide excitation or inhibition to similarly tuned neurons in the contralateral ICc, while NPY commissural neurons in the ICc are more likely to provide cross-frequency lateral inhibition to the contralateral ICc. The ICc injection cases also showed that both NPY and non-NPY neurons in the ICd send a large projection to the contralateral ICc. The commissural projection from the ICd to the ICc lacked any clear differences between non-NPY and NPY neurons. Commissural projections from the IClc were much less common and, when present, also lacked a clear organization.

### Commissural projections to the ICd primarily originate from the contralateral ICd

In one mouse, Case 36, the RB injection was located entirely within the right ICd (**Figure 6A**). Previous studies report that commissural projections to the ICd largely arise from locations distributed throughout the contralateral ICd (i.e., a heterotopic, divergent projection) and to a lesser extent from regions of the contralateral ICc that have similar frequency tuning to the ICd region under study (i.e., a tonotopic projection; Saldaña and Merchán, 1992; Malmierca et al., 2009). Consistent with the former observation, most of the RB-labeled NPY and non-NPY neurons in the left IC of Case 36 were in the ICd, with no clear organization within the ICd other than a tendency to be denser near the site homotopic to the injection site (**Figure 6B**). A smaller number of RB-labeled NPY and non-NPY neurons were observed in the left ICc, along with an even smaller number in the left IClc, but there was no clear organization to these distributions. Heatmaps of the density of RB-labeled neurons located in the ICd (ICc and IClc neurons removed) revealed a different pattern than observed with the ICc injection sites: RB-labeling for both non-NPY neurons and NPY neurons largely overlapped and was densest at the site homotopic to the ICd injection site (**Figure 6C**). Analysis of the distances of RB-labeled ICd neurons to the site homotopic to the centroid of the RB injection site did not detect a significant difference between the distribution of NPY and non-NPY neurons (**Figure 6D**; medians, NPY = 304 µm, non-NPY = 305 µm; two-sample Kolmogorov-Smirnov test: *n* = 319 non-NPY, 73 NPY, *D*_(319,73)_ = 0.11, *p* = 0.43). Similar results were obtained when this analysis was repeated using only the two-dimensional distances of neurons to the homotopic centroid (rostral-caudal dimension discarded; medians, NPY = 297 µm, non-NPY = 291 µm; two-sample Kolmogorov-Smirnov test: *D*_(319,73)_ = 0.11, *p* = 0.47).

**Figure 6.**
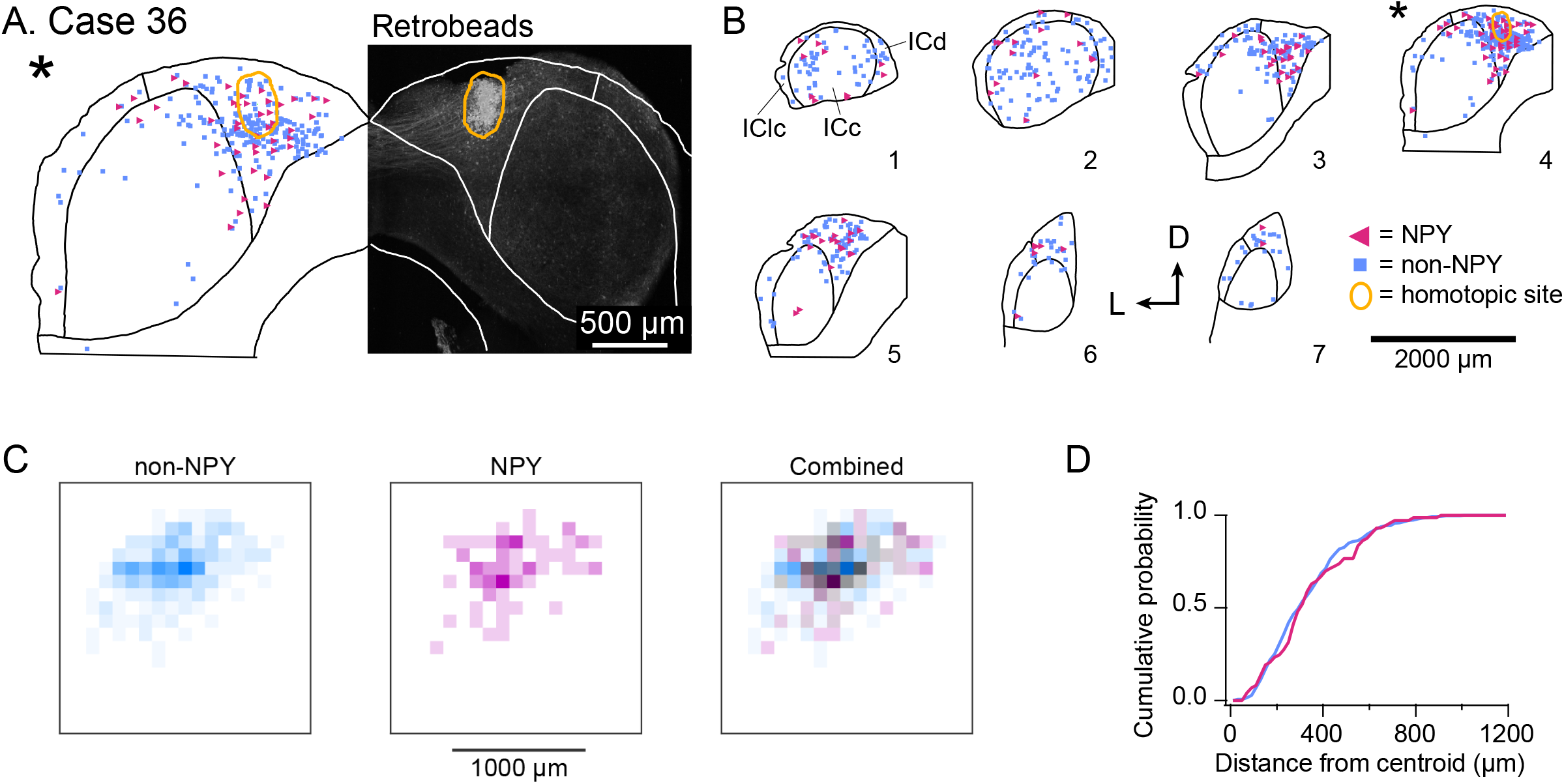
NPY and non-NPY neurons in the ICd have similar commissural projection patterns. Results from Case 36 where the RB injection was constrained to the right ICd. **A** and **B** show the tracer deposit and distributions of labeled cells as in Figure 3. **A)** RB deposit site in the right ICd and homotopic site in the left IC in experimental Case 36. The gray-scale image shows the center of the RB deposit (circled in gold). The left half of the panel shows an outline of the left IC with the homotopic location of the injection site indicated by the gold outline. The illustrated section (*) is from a series of sections shown in **B. B)** The full series of plots for Case 36 showing the distributions of RB-labeled (commissural) neurons that were NPY+ (magenta triangles, *n* = 119 neurons) and non-NPY (cyan squares, *n* = 598 neurons). Sections are numbered from caudal to rostral. The asterisk indicates the section containing the injection site shown in **A**, and the area homotopic to the injection is outlined in gold. RB-labeled neurons were distributed throughout the ICd, including at the site homotopic to the injection site. D – dorsal; L – lateral. **C)** Heatmaps show the densities of non-NPY commissural neurons (cyan, left) and NPY commissural neurons (magenta, middle) in 100 µm^2^ bins in the left ICd. In contrast to the ICc injection cases, a merged overlay of the non-NPY and NPY densities (right) reveals that RB-labeled NPY and non-NPY neurons had largely overlapping distributions in the ICd. **D)** A cumulative probability plot shows the distributions of the distances of RB-labeled non-NPY (cyan) and NPY (magenta) neurons in the left ICd from a point homotopic to the centroid of the RB injection site in the right ICd. A two-sample Kolmogorov-Smirnov test revealed that the distributions of distances to the centroid did not significantly differ for NPY and non-NPY neurons (see Results for details). ICc and IClc neurons were excluded from the analyses in **C**,**D**.

As these data come from one injection case, they are not sufficient to support strong conclusions, but we include the results here because they are consistent with past studies and bolster support for the hypothesis that commissural projections targeting the ICd primarily originate in the contralateral ICd. Furthermore, the lack of a clear difference in the distribution of NPY and non-NPY commissural neurons targeting the contralateral ICd contrasts with the differences observed for NPY and non-NPY commissural neurons targeting the contralateral ICc. Together, these results suggest that the organizing principles and corresponding functional roles of commissural projections differ depending on the IC subdivision that is targeted and the neuron type forming the projection.

## Discussion

We found that NPY neurons provide ∼11% of the commissural projection in the IC, likely accounting for a large portion of the inhibitory commissural projection. Within the ICc, the commissural projections of NPY neurons were more heterotopic and more likely to connect regions with different frequency tuning than the commissural projections of non-NPY neurons (**Figure 7A**). This pattern appeared exclusive to the ICc; commissural projections targeting the ICd were similarly organized whether they came from NPY neurons or non-NPY neurons (**Figure 7B**). The commissural projections of non-NPY neurons largely matched patterns reported in previous studies, connecting homotopic isofrequency bands in the ICc and exhibiting a more dispersed connection pattern in the ICd. Together, our results indicate that a large portion of the inhibitory commissural pathway follows a different organizing logic than that followed by most commissural neurons. Based on the tonotopic organization of the ICc, we propose that commissural NPY neurons in the ICc play a special role in bilateral integration by providing cross-frequency lateral inhibition to neurons in the contralateral ICc.

**Figure 7.**
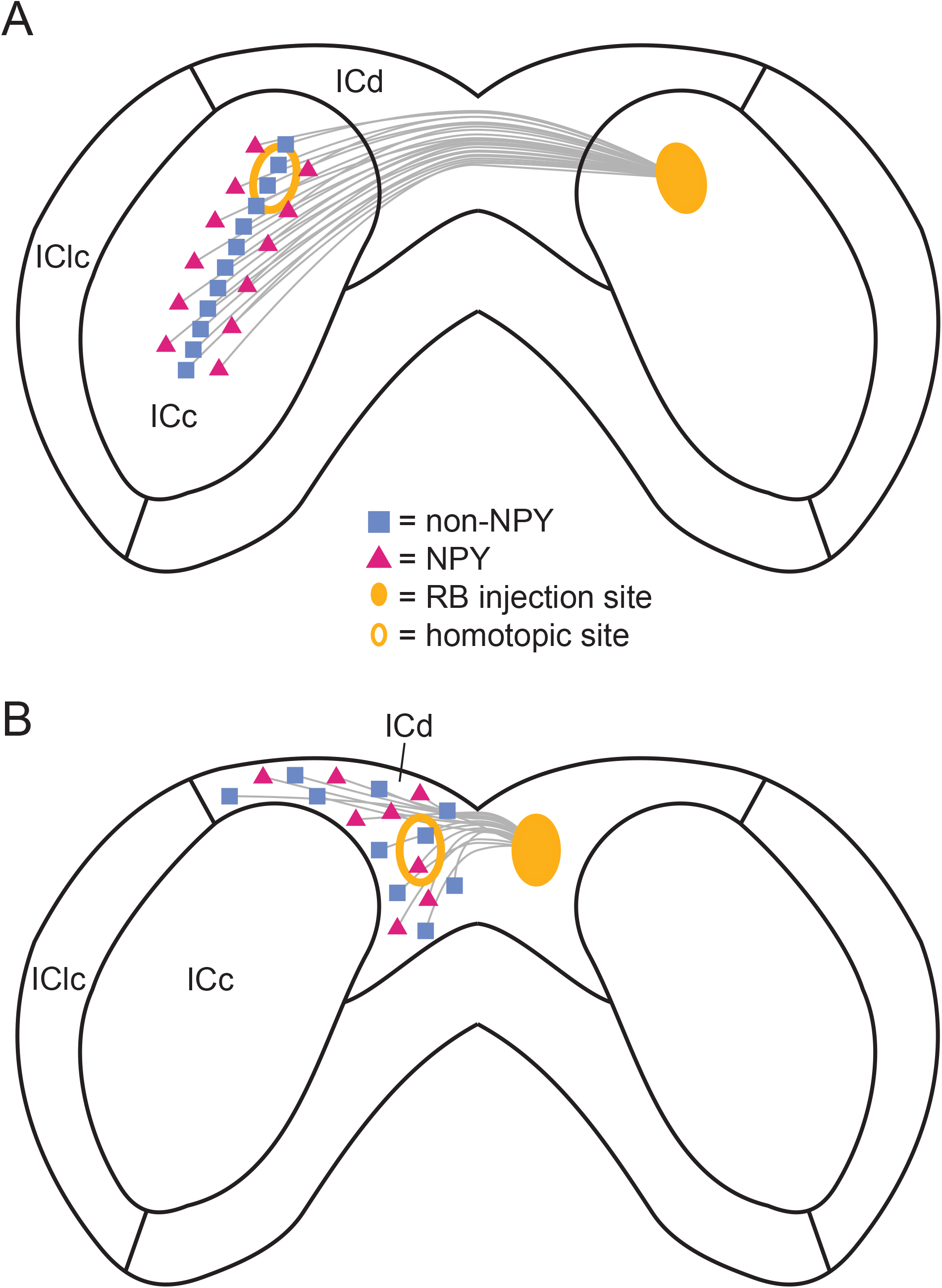
Summary of the commissural projection patterns of NPY and non-NPY neurons in the ICc and ICd. **A)** Commissural projections between the left and right ICc. The commissural projections of non-NPY neurons tended to be homotopic and to connect similarly tuned frequency laminae in the left and right ICc. In contrast, the commissural projections of NPY neurons were more heterotopic and more likely to arise from regions bordering the region where non-NPY commissural neurons were found. These patterns suggest that non-NPY neurons connect regions of the left and right ICc that have similar frequency tuning, while NPY neurons connect regions with dissimilar frequency tuning, possibly providing a source of sideband inhibition to the contralateral ICc. **B)** Commissural projections between the left and right ICd. The commissural projections of non-NPY and NPY neurons in the ICd were similarly heterotopic, suggesting that commissural projections between the left and right ICd follow a different pattern than those joining the left and right ICc.

### The inhibitory commissural projection

Our data show that NPY neurons account for ∼11% of commissural neurons in the IC, indicating that at least 11% of the IC commissural projection is inhibitory. Past studies have disagreed about the inhibitory contribution to the commissural projection. The first studies to use double-labeling to identify GABAergic commissural neurons had conflicting results, one reporting that ∼37-39% of the IC commissural projection in rats was inhibitory (González-Hernández et al., 1996) while the other reported no GABAergic contribution to the commissural projection in rats (Zhang et al., 1998). Interestingly, González-Hernández et al. observed five-times more labeling of commissural GABAergic neurons when tracer injections were made in the dorsal ICc compared to when injections were made in the ventral ICc. Consistent with this, our ICc injection cases tended to show more labeling of NPY neurons in the dorsal region of the contralateral ICc than the ventral region, even when the injection site was in the ventral ICc (Cases 22, 08). These results support the hypothesis that GABAergic commissural neurons are more common in the dorsal ICc. Similarly, the apparent absence of commissural GABAergic neurons in the Zhang et al. study from 1998 may have resulted from tracer injections localized to the ventral ICc. A more recent study in rats reported that ∼20% of ICc commissural neurons were GABAergic, but the sample size included only six GABAergic neurons (Hernández et al., 2006). In contrast, a larger study in guinea pigs found that ∼9% of the commissural projection was GABAergic (Nakamoto et al., 2013). In mice, a large screening study found that ∼34-38% of commissural inputs to excitatory IC neurons were GABAergic, while ∼22% of commissural inputs to ICc inhibitory neurons and 65% of commissural inputs to inhibitory neurons in the IC shell (ICd + IClc) were GABAergic (Chen et al., 2018).

Discrepancies in estimates of the size of the inhibitory commissural projection likely come from methodological differences, technical limitations, and species differences. However, the wealth of evidence strongly supports the conclusion that GABAergic neurons make a significant contribution to the IC commissural projection. Since NPY neurons account for approximately one-third of IC GABAergic neurons (Silveira et al., 2020), our finding that NPY neurons comprise ∼11% of commissural neurons likely provides a lower bound for the total contribution of inhibitory neurons to the commissure. This result would be consistent with the higher percentages of GABAergic commissural neurons reported by most past studies in rodents.

Interestingly, while anatomical surveys indicate that a minority of commissural neurons are inhibitory, inhibitory synaptic input is commonly observed when the commissure is electrically stimulated in brain slices (Smith, 1992; Moore et al., 1998; Li et al., 1999; Reetz and Ehret, 1999). Similarly, using optogenetics to activate commissural input to VIP neurons in the IC, we found that inhibitory postsynaptic potentials were a more common occurrence than excitatory postsynaptic potentials (50% of neurons compared to 41% of neurons, respectively)(Goyer et al., 2019). Based on our present results, a possible explanation for these observations is that inhibitory commissural neurons might have more divergent axons than excitatory commissural neurons. Instead of the homotopic projections that typified non-NPY commissural projections to the ICc, the distribution of commissural NPY neurons was more dispersed, possibly indicating a larger degree of branching and more postsynaptic targets for NPY axons. Future studies of the morphology of commissural axons will be key to testing this idea.

### Homotopic versus heterotopic commissural projections

One of the most consistent observations about the organization of the IC commissural projection is that commissural inputs to ICc neurons are homotopically and tonotopically organized (González Hernández et al., 1986; Saldaña and Merchán, 1992; Malmierca et al., 1995, 2009). Our data on the commissural projections of non-NPY neurons fit this pattern, with a clear tendency of non-NPY commissural neurons to be densest at the site homotopic to the RB injection site and, for RB injections in the ICc, to spread along a dorsomedial to ventrolateral axis consistent with the shape of the isofrequency lamina in the ICc (cf. Meininger et al., 1986). Surprisingly, commissural NPY neurons in the ICc broke from this trend, often being at low density or even absent from the site homotopic to the RB injection and more often located lateral and medial to the main band of non-NPY neuron labeling. Due to the tonotopic organization of the ICc, we interpret this result to mean that commissural NPY neurons in the ICc are more likely to target neurons in the contralateral ICc located in isofrequency bands adjacent to the homotopic isofrequency band. This points to a possible role for commissural NPY neurons in providing a contralateral source of cross-frequency lateral inhibition (i.e., what is often referred to as sideband inhibition).

An important question is whether the organization of the NPY commissural projection is unique to NPY neurons or a feature shared with other inhibitory neurons or other stellate neurons. As mentioned above, NPY neurons comprise approximately one-third of IC GABAergic neurons, and it therefore seems reasonable to predict that other classes of IC GABAergic neurons also contribute to the IC commissure. In addition, past studies have indicated that both disc-shaped and stellate neurons in the ICc contribute to the commissural projection (González Hernández et al., 1986; Okoyama et al., 2006; Nakamoto et al., 2013; Goyer et al., 2019), and it is known that disc-shaped and stellate neurons come in both glutamatergic and GABAergic varieties (Oliver et al., 1994; Ono et al., 2005b). Since stellate neurons comprise a minority of ICc neurons and their dendritic morphology suggests that they typically integrate input from two or more isofrequency lamina, it is appealing to hypothesize that disc-shaped neurons comprise the major, homotopic portion of the ICc commissural projection while ICc stellate neurons provide the minor, heterotopic commissural projection. NPY neurons represent approximately 38%-50% of ICc stellate neurons (Silveira et al., 2020), and therefore it will be important to compare the projection patterns of NPY neurons to other classes of commissural stellate neurons, such as VIP neurons, which comprise another ∼18%-23% of stellate neurons (Goyer et al., 2019).

In contrast, the organization of the commissural projection to the ICd was similar for NPY and non-NPY neurons, with the densest projection originating from a homotopic site and a large remaining projection originating from dispersed sites throughout the ICd. Since these data were from a single case it is not clear yet whether these are representative results, but it is striking that homotopic labeling was minimal for NPY neurons in the ICc injection cases but was strong in the ICd case. Our results are consistent with the past observations that the commissural projection to the ICd is more dispersed than that to the ICc (Malmierca et al., 2009) but also indicate that there may be some tendency for ICd commissural projections to connect homotopic ICd regions. Overall, the number of ICd injection cases in the present study and past studies are very few, pointing to a clear need for a more in-depth examination of the ICd commissural projection in the future.

### Functional role of the NPY commissural projection

Previous studies point to a modulatory role for the IC commissural pathway, wherein commissural input adjusts but does not set the receptive fields of neurons in the contralateral IC (Malmierca et al., 2003, 2005; Orton and Rees, 2014; Orton et al., 2016). In these studies, blocking commissural input typically changed the gain of the input-output functions describing how IC neurons altered their firing in response to changing auditory cues. Since commissural gain control generally enhanced the dynamic range and sensitivity of IC neurons to tones (Malmierca et al., 2003, 2005; Orton and Rees, 2014) and azimuthal sound localization cues (Orton et al., 2016), it appears that commissural input may boost the capacity of IC neurons to detect cues close to the detection threshold and/or to discriminate among related auditory cues.

Because past functional studies employed techniques that broadly blocked neurons in the contralateral IC, it is unclear how inhibitory and excitatory commissural projections separately contribute to these effects, and it is completely unknown how individual classes of commissural neurons shape coding in the contralateral IC. Since NPY neurons are molecularly identifiable, it should now be possible to test functional hypotheses about how NPY commissural neurons shape sound coding. For example, our results suggest that NPY commissural neurons provide cross-frequency lateral inhibition to neurons in the contralateral IC, and this could be directly tested by comparing the width of the frequency response areas of IC neurons under control conditions and when NPY neurons in the contralateral IC are inhibited via optogenetic or chemogenetic approaches. We predict that frequency response areas would show stronger sideband inhibition under control conditions than when NPY neurons in the contralateral IC are inhibited. Such approaches with NPY neurons and other identified classes of commissural neurons will be key to uncovering the circuit logic and computations underlying commissural modulation in the IC.

## Acknowledgements

This work was supported by a Magnificent Michigan Fellowship from the Kavli Neuroscience Innovators at the University of Michigan (JDA) and by National Institutes of Health Grants R01 DC018284 (MTR) and R01 DC004391 (BRS). Special thanks to Nina Lenkey for technical assistance.

